# Stabilizing selection shapes variation in phenotypic plasticity

**DOI:** 10.1101/2021.07.29.454146

**Authors:** Dörthe Becker, Karen Barnard-Kubow, Robert Porter, Austin Edwards, Erin Voss, Andrew P. Beckerman, Alan O. Bergland

**Affiliations:** Department of Biology; University of Virginia, Charlottesville/VA, USA; Department of Animal and Plant Sciences; University of Sheffield, Sheffield, UK; Department of Biology, James Madison University, Harrisonburg/VA, USA; Biological Imaging Development CoLab, University of California San Francisco, San Francisco/CA, USA; Department of Integrative Biology, University of California Berkeley, Berkeley/CA

**Keywords:** stabilizing selection, phenotypic plasticity, heritability, *Daphnia*, defense morphologies

## Abstract

The adaptive nature of phenotypic plasticity is widely documented in natural populations. However, little is known about the evolutionary forces that shape genetic variation in plasticity within populations. Here we empirically address this issue by testing the hypothesis that stabilizing selection shapes genetic variation in the anti-predator developmental plasticity of *Daphnia pulex*. The anti-predator morphological defense is characterized by pedestal and spikes that grow on the back of the *Daphnia* neck following exposure to predator cure. We characterized variation in this plasticity using a novel, high-throughput phenotyping method that describes the entire dorsal shape amongst >100 *D. pulex* strains originating from a natural population in the UK. We found low genetic diversity for morphological defenses among genetically diverse clones upon predation risk exposure. The strongest reduction in genetic variation was observed in areas of greatest phenotypic plasticity, which we interpret as evidence of stabilizing selection. By assessing among-clone variance in clonally related, field derived strains, we contrasted mutational variation (V_m_) to standing variation (V_g_). We found that V_g_/V_m_ is lowest in areas of greatest plasticity. These data strongly suggest that stabilizing selection operates directly on phenotypic plasticity, providing a rare glimpse into the evolution of fitness related traits in natural populations.

## Introduction

Organisms frequently experience temporal and spatial fluctuations in their natural habitats. The capacity to persist and thrive across variable environmental conditions is due to local adaptation at the population level, and phenotypic plasticity at the individual level. Phenotypic plasticity, the ability of a single genotype to alter its phenotype in response to environmental change, is ubiquitous in natural populations and well established as an adaptive response to environmental change [1, 2]. Providing time for a population to become established, adaptive plasticity reduces the probability of extinction in new or fluctuating environments [3, 4], and also enables populations to efficiently traverse fitness landscapes [3]. Decades of theoretical and empirical work on the quantitative genetics of plasticity has revealed that genetic variation for plasticity exists [5, 6], that heritable variation in plasticity can respond to natural selection [1, 7] and that plasticity is itself locally adapted (e.g., [8]).

Evidence for the adaptive nature of phenotypic plasticity has been demonstrated for morphological and life-history traits [9]. Broadly, this work is aligned with theoretical predictions that adaptive phenotypic plasticity evolves when environmental change is rapid [10, 11], environmental cues are reliable [12, 13], plastic responses occur as rapidly as environmental change [14], and when the incurred cost of plasticity is low [15, 16]. Empirical studies on the adaptive nature of phenotypic plasticity often measure aspects of fitness following environmental exposures (e.g., [17]), or document the plastic responses of genotypes collected from different locations that reflect alternate historical selection regimes (e.g., [18]).

However, such data and insights do not reveal the selective forces shaping levels of genetically based phenotypic variation within populations. Theoretical models aimed at explaining the level of quantitative genetic variation in natural populations frequently highlight the central role of stabilizing selection [19, 20]. Stabilizing selection reduces functional genetic variation in a population without modifying the population mean and maintains populations near their local fitness peak. Yet there is a stark gap between the theoretical expectation that stabilizing selection commonly operates on natural populations and the empirical evaluation of stabilizing selection [21, 22] (but see [23, 24]). This data deficit is especially pronounced for phenotypic plasticity, because of the compounding challenges of measuring genetic variation in plasticity (e.g., [12, 25]) and of estimating mutational variation, which is required to test hypotheses about the mechanisms that shape the magnitude of genetically based phenotypic diversity [25, 26].

We directly address this gap by evaluating the signature of stabilizing selection on predator induced defense in the eco-evolutionary model, *Daphnia pulex*. This species can develop a defensive, spikey pedestal when in the presence of predatory *Chaoborus spp.* larvae and is a century old, classic example of adaptive phenotypic plasticity [27, 28]. The predator induced defenses are size and age specific and lead to an increased survival to predator attack by up to 50% [28, 29]. Induction in the absence of actual predators incurs a fitness cost [28, 30].

We combine a novel, high-throughput phenotyping assay with genome re-sequencing to test the hypothesis that this plastic response is subject to stabilizing selection. We first show that genetic variation for the defense, in an outbred population is reduced upon exposure to predator cue. The reduction of genetic variation is greatest in the region of the dorsal axis with greatest plasticity. We demonstrate that this region is an independent phenotypic module, suggesting that the pleiotropic costs to plasticity may have been minimized. To evaluate the signature of stabilizing selection, we measured mutational variation of plasticity using clonally related strains, and contrasted mutational variation (V_m_) to standing variation (V_g_) in the outbred individuals. Consistent with the view that plasticity is subject to stabilizing selection, we show that V_g_/V_m_ is lowest in the region of greatest plasticity. Taken together, these data provide unique insight into the evolution of phenotypic plasticity and the forces that shape genetic variation in the wild.

## Results

### Robust and accurate phenotyping

To date, assessment of predator induced morphological changes in *Daphnia* have been based on a categorical scoring technique that classifies pedestals as absent (score = 0), small (score = 30) or large (score = 50) and adds spikes in increments of 10 [28, 29]. Although this scoring technique has been useful, its coarse scale and potential for observer bias limits its capacity to assess plasticity with high resolution, high replication and reproducibility, and low observer bias.

To remedy this, we developed a novel analysis tool, *DAPCHA*, that allows automated identification of defined phenotypic landmarks and subsequent quantification of phenotypic plasticity using standardized photographic images from stereoscope microscopy (Fig. 1A). These features include basic aspects of morphology such as animal length, eye size, tail length (Fig. 1A-II), as well as a trace of the entire dorsal edge (Fig. 1A-III). *DAPCHA* accurately measures animal length (Fig. 1B; Suppl. Tab. 1) and is more reliable in unit conversion of length estimates than are manual observers (e.g., run 2, Fig. 1C; Suppl. Tab. 1).

**Figure 1.**
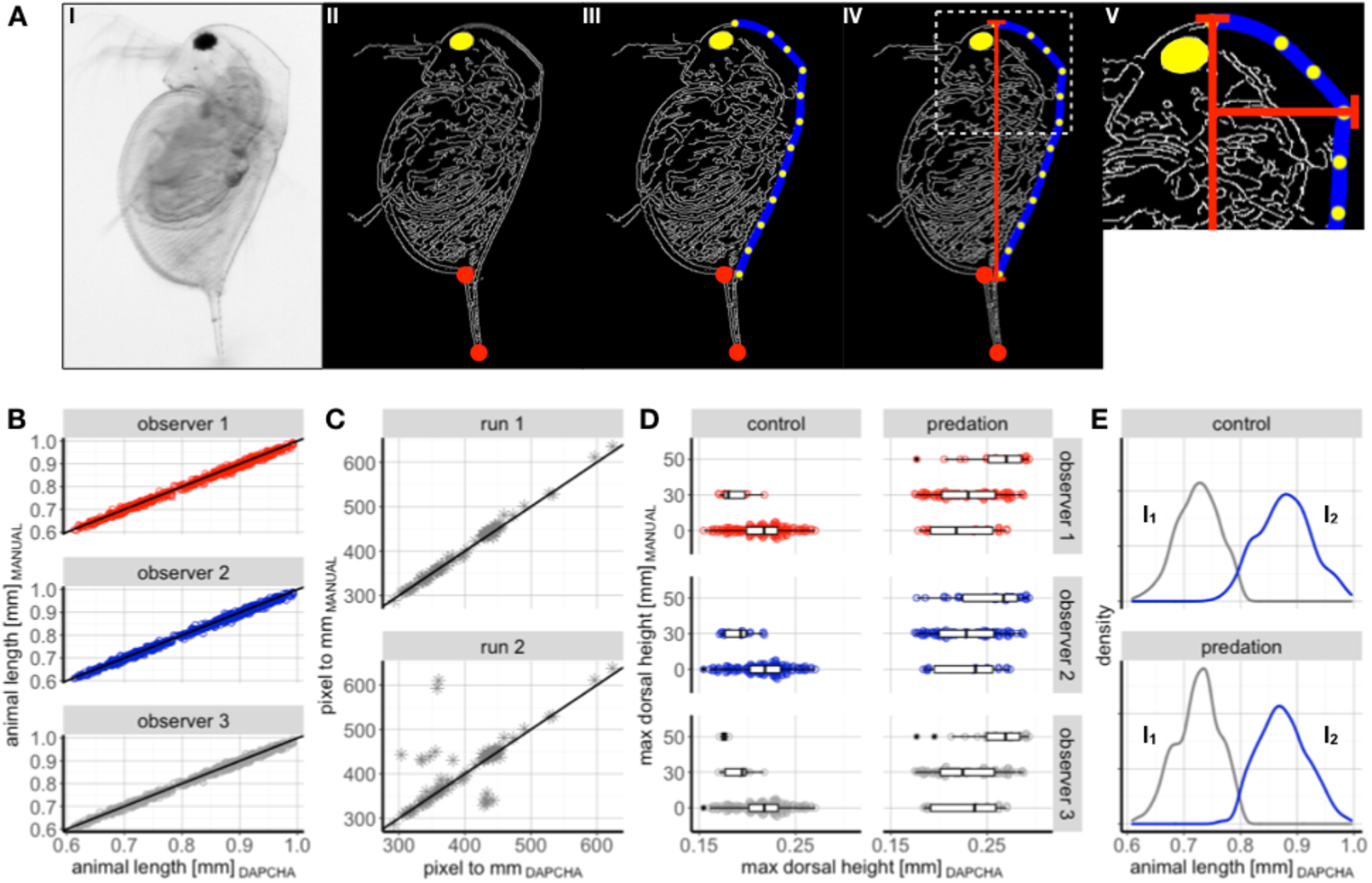
High-throughput phenotypic assessment via novel, automated image analysis tool ‘*DAPCHA’*. (A) Phenotypic assessment of *Daphnia* using *DAPCHA* involves three major steps: conversion of a standardized, raw image to greyscale (I-II); automated identification of key landmarks (i.e., eye, tail tip and tail base) (II); and automated tracing of the dorsal edge of the carapace (blue line) via identification of equally spaced landmarks along the dorsal axis (yellow points) (III). Defined landmarks subsequently allow for the quantification of different phenotypic traits, including animal length (IV) and dorsal height (V; here exemplified by the dorsal position where dorsal height was largest). (B-D) Accuracy of phenotypic estimates by *DAPCHA* were validated via contrasting manual estimates with automated data: animal length estimates across three different observers (B), unit conversion of length estimates using a microstage meter assessed in two different runs (see Materials and Methods) (C), and morphological changes in the nuchal area under control and predation conditions estimated by three different observers (D). (E) Using a mixture model on animal length and animal area, test animals were retroactively assigned to distinct developmental stages (i.e., first instar, I1; second instar, I2).

We used *DAPCHA* to calculate the maximum dorsal height of the animal (Fig. 1A-V, and see Materials and Methods), as a summary of morphological plasticity before and after exposure to predator derived kairomones. We find that maximum dorsal height significantly changes with the production of pedestals and spikes in response to predator cue (Fig. 1D; Suppl. Tab. 1) and is also strongly correlated with manual assessments based on the discrete manual scale, described above (Suppl. Tab. 1). We also show that there was inconsistent manual assessment of the pedestal score between three independent observers (Fig. 1D; Suppl. Tab. 1), suggesting that *DAPCHA* is a useful approach to assess plasticity in an unbiased manner. We retroactively assigned animals to the first instar and second instar (Fig. 1E) based on the multimodal distribution of animal length and animal dorsal area (see Materials and Methods), enabling us to ask questions about the extent of plasticity in distinct size and age classes.

### Evidence for stabilizing selection in an outbred sample

We estimated components of phenotypic variation in the plastic response among 49 genetically unique strains (Suppl. Fig. 1) sampled from a single population in southern England ([31]). We reared each strain with and without kairomone and photographed ca. 4 individuals per strain and treatment group each day, for the first 3-4 days post-parturition (471 individuals, Suppl. Fig. 2).

First, we evaluated the basic plastic response of these strains. We demonstrate that exposure to kairomone induces a plastic response, most prominently in the nuchal area (Fig. 2A,E; Suppl. Fig. 3). The magnitude and position of the defense varied with predation risk. The maximum dorsal height shifted towards anterior head regions (Fig. 2A,B; Suppl. Fig. 3; Suppl. Tab. 1) and the maximum height of the defense structures increased (Fig. 2C; Suppl. Tab. 1). Furthermore, we detected an increase in the number of neckteeth under predation risk (Fig. 2D; Suppl. Tab. 1). A quantitative genetic test of modularity (see Materials and Methods) indicates that standing genetic variation in the region of maximum plasticity is uncorrelated with the rest of the *anterior-posterior* (A-P) axis, suggesting that this region responds to selection as an independent phenotypic module (Suppl. Fig. 4). Such a pattern suggests that pleiotropic costs to plasticity may have been minimized.

**Figure 2.**
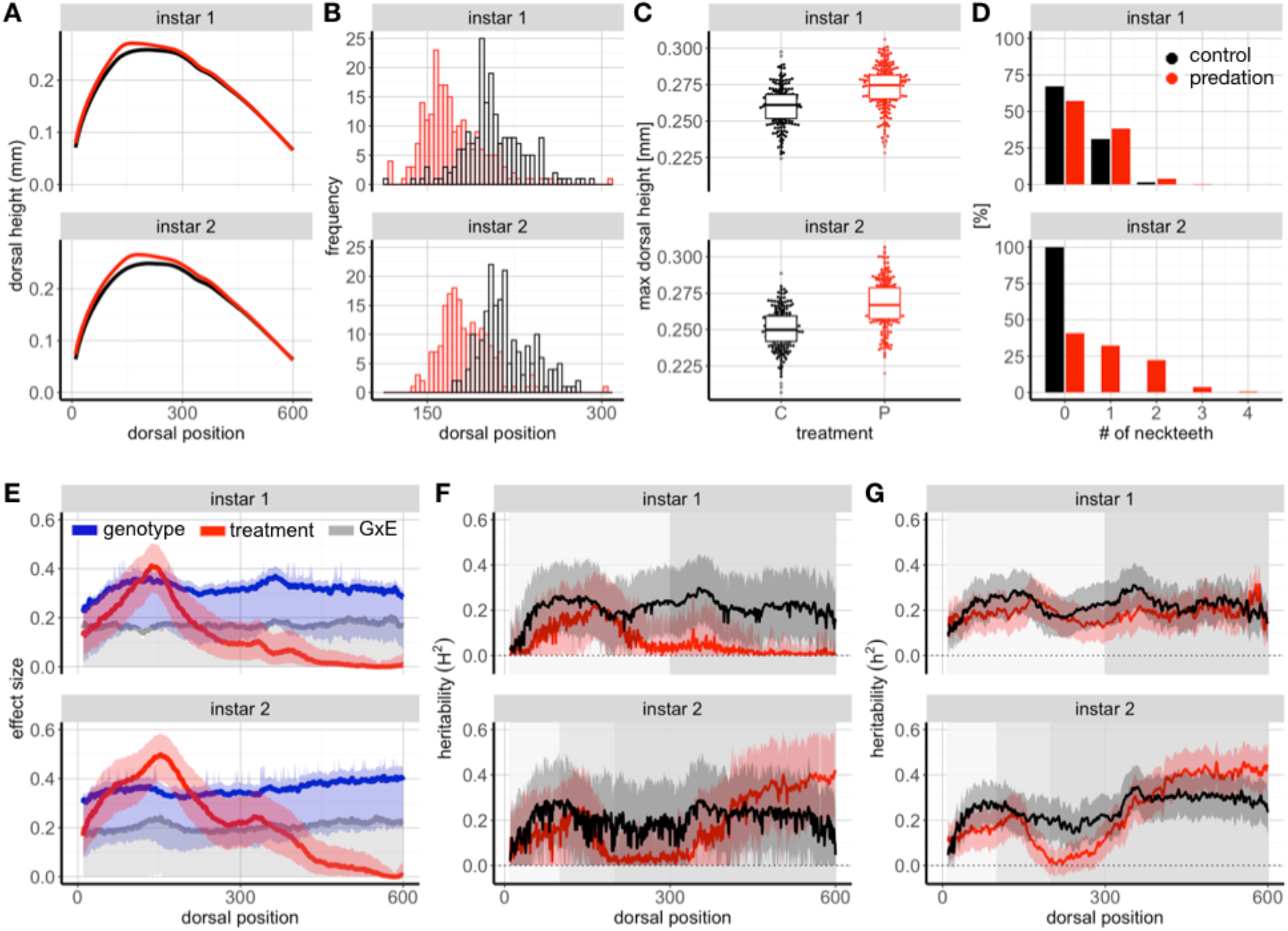
Effects of predation risk on morphological changes in genetically unique strains. (A) Risk of predation induces marked shape changes along the dorsal axis in first and second instar *D. pulex* (control: black line; predation: red line). (B) Strongest morphological changes are observed in the head area, with maximal dorsal height shifting towards anterior head regions under predation risk. (C) Predator induced defense morphologies, here measured as maximal dorsal height, increase in response to predation risk exposure in both instars (control, C: black points; predation, P: red points). (D) The number of neckteeth increases in response to predation risk exposure, particularly in second instar animals. (E) Effect sizes from a linear model along the dorsal axis reveal distinct patterns of treatment (i.e., predation risk, red line), genotype (blue line), and GxE (grey line) effects on morphological changes. Shaded areas indicate upper (0.95) and lower (0.05) confidence intervals. (F,G) Broad sense (F) and narrow sense (G) heritability estimates of dorsal height vary along the dorsal axis in response to control conditions (black line) and predation risk (red line), with a strong reduction of both measures of heritability for dorsal height upon predation risk exposure in the region of maximum plasticity (i.e., dorsal positions 100-250). Shaded areas indicate upper (0.95) and lower (0.05) confidence intervals (F) or standard errors (G). Grey rectangles in panels F and G highlight morphological independent shape modules, separating head and posterior body areas (see Suppl. Fig. 4).

We decomposed phenotypic variation in plasticity among these individuals to estimate the relative contribution of genotype, environment, and genotype-by-environment at each position along the dorsal A-P axis. We observe a strong plastic response of dorsal height among our outbred population (Fig. 2E, red line) and, as expected, the peak of induction is centered in the nuchal area around dorsal position 150. We observed substantial genetic variation in the induced morphological defense (Fig. 2E, blue line).

There is a pronounced increase in GxE variation for instar 2 around position 150 (Fig. 2E, grey line). This elevated GxE near the region of maximum induction could be caused by crossing reaction norms or by a change in genetic variance between the control and predator cue environments. To test these alternative explanations, we calculated both the broad- and narrow-sense heritability of dorsal height in the two environments. We find a reduction of both measures of heritability for dorsal height upon exposure to predator cue slightly dorsal to the region of maximum plasticity (Fig. 2F,G; Suppl. Tab. 1). This result is consistent with the hypothesis that phenotypic plasticity is subject to stabilizing selection.

### Mutational variation and further evidence of stabilizing selection

We contrasted levels of standing genetic variation, V_g_, with mutational variation, V_m_, as an additional test for stabilizing selection [32]. To estimate mutational variation in phenotypic plasticity, we characterized 56 clonally related strains (Suppl. Fig. 1) using the same experimental design as described above (516 individuals) and assayed concurrently with the genetically unique clones. These clonally related strains were independently isolated from the field and sampled from the same ponds at the same time as the outbred individuals, above, allowing us to directly relate levels of genetic variation in the plastic response. Population genomic analysis suggests that these clonally related isolates share a recent common ancestor and are also related to the outbred individuals that we studied here ([31]). Detailed assessment of the plastic response of these clonally related strains and their among-line variance reveal that these clonally related strains display a robust plastic response that is similar to the average plastic response of the outbred individuals (Suppl. Fig. 5). We also detected considerable variation in plasticity among these clonally related strains in several key metrics of plasticity such as the height along the A-P axis (Suppl. Fig. 5A; Suppl. Tab. 1), and the maximum height of the pedestal (Suppl. Fig. 5C; Suppl. Tab. 1).

We confirmed that the phenotypic variance that we observe among these clonally related strains is heritable, and unlikely to be caused by other factors such as maternal effects or experimental artifact, by performing a ‘twin analysis’. For this analysis, we took advantage of a feature of our experimental design, evaluating the correlation of plasticity between siblings released from the same mother (‘within clutch’), and between individuals from the same strain but born to different mothers (‘within clone’), replicated across the 56 strains of the same clonal assemblage. If the significant among-line variance that we observe across the clonally related strains were due to experimental artifact or shared environment, the correlation between individuals released from the same mother (‘twins’; within clutch) should be higher than the correlation between individuals released from different mothers of the same strain (‘cousins’; within clone). Similarly, if the among-line variance of the clonally related strains is due experimental artifact, the correlation between individuals within a batch should be high. On the other hand, if new mutations (or heritable epigenetic marks, see e.g., [33]) cause phenotypic differentiation between clonally related strains, then twins should be (i) as similar to each other as cousins, and (ii) should be more similar to each other than to other genetically similar lineage of the clonal assemblage (among clones).

In line with the expectation that mutations underlie variation, we detected strong correlation between twins and between cousins and that these individuals are more similar to each other than to randomly paired individuals sampled across the clonal assemblage (Fig. 3A; Suppl. Tab. 1). Moreover, the correlation coefficients between twins broadly exceeded the expectations from permutations (Fig. 3A, ‘within clutch’; Suppl. Tab. 1), while correlations of randomly paired individuals from the clonal assemblage were markedly lower (Fig. 3A, ‘within clutch’ and ‘among clones’; Suppl. Tab. 1). We also detected no considerable correlation of phenotypic responses among individuals from the same experimental batches (Fig. 3A, ‘among batches’; Suppl. Tab. 1), suggesting that both experimental batch and maternal effects are not driving the observed variance among clonally related strains.

**Figure 3.**
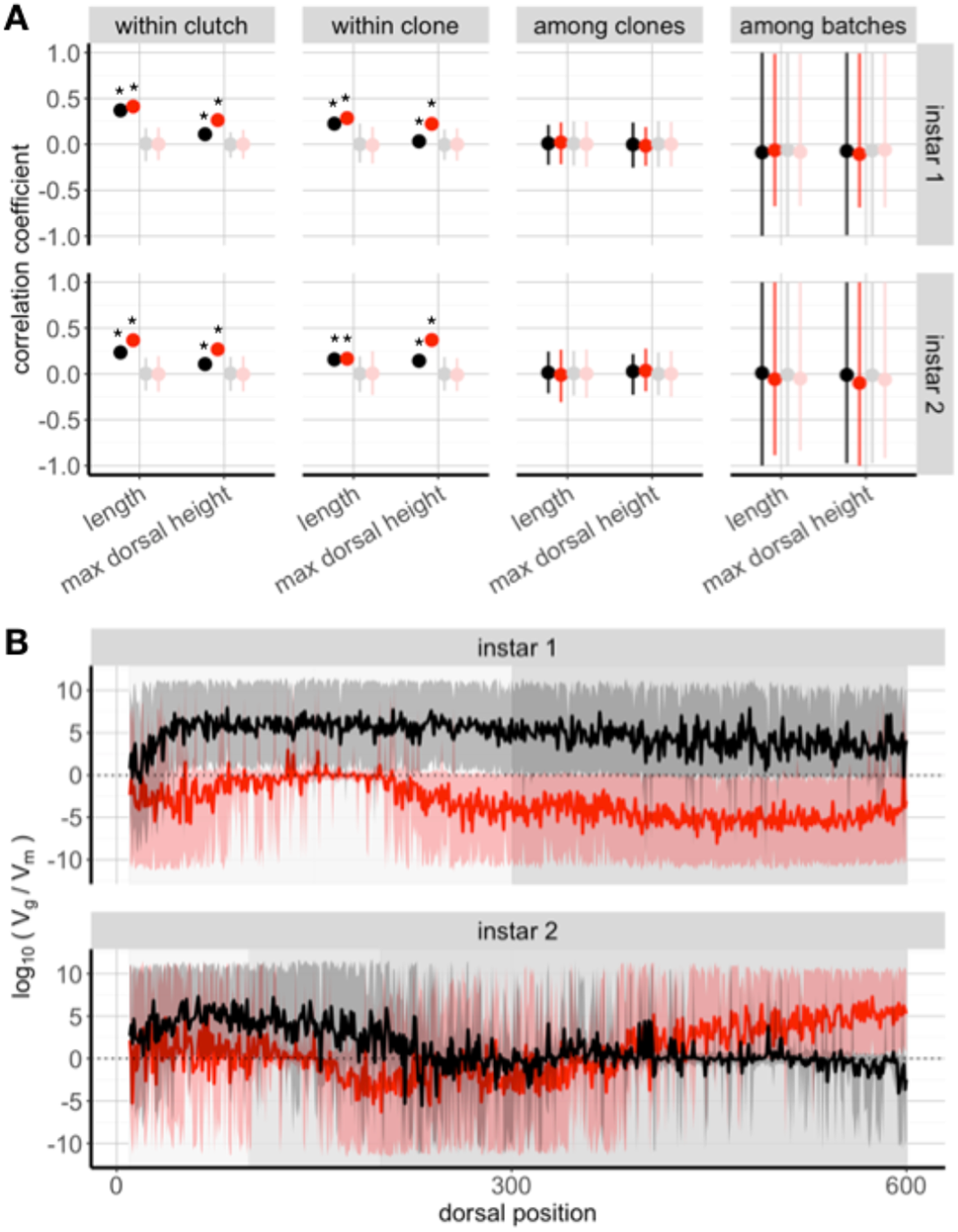
Effects of natural selection on predator induced plasticity in *Daphnia*. (A) Phenotypic variation in genetically similar strains arises due to genetic effects: phenotypic responses (here: animal length and maximal dorsal height) of offspring released from the same mother (‘within clutch’) and same strain (‘within clone’) are more similar to each other than to offspring released from a randomly drawn member of the clonal assemblage (‘among clones’). Correlation coefficients broadly exceed coefficients calculated for permuted data. Moreover, phenotypic correlations among randomly paired individuals from the same experimental batches are low, with actual data not exceeding permuted data ranges. Black and red points indicate control and predation risk conditions with darker and lighter colors depicting actual and permuted data, respectively. Error bars indicate upper (0.95) and lower (0.05) confidence intervals. Asterisks indicate actual data exceeding permuted data ranges (see Suppl. Tab. 1). (B) log10(Vg/Vm) estimates of dorsal height vary along the dorsal axis in response to control conditions (black line) and predation risk (red line). Notably, in second instar *Daphnia*, genetic diversity is strongly reduced in areas of largest phenotypic plasticity. Shaded areas indicate upper (0.95) and lower (0.05) confidence intervals. Grey rectangles highlight morphological independent shape modules, separating head and posterior body areas (see Suppl. Fig. 4).

From this analysis, plus the linear model approach described above, we conclude that there are heritable differences in plasticity between strains that are clonally related. We interpret this among-line variance as mutational variance, V_m_. The magnitude of V_m_ that we document here is on par with estimates of mutational variance in other *Daphnia* studies [34]. The large mutational variance also demonstrates that plasticity is not mutation limited and suggests that local adaptation in the plastic response (e.g., [8]) could, in principle, be driven by new mutations that arise frequently.

Under a model of mutation-selection balance, which assumes that mutations are either neutral or mildly deleterious, the strength of stabilizing selection on quantitative traits can be estimated from the ratio of standing genetic variation and the mutational variance [35]. We used the among line variance of genetically unique strains as an estimate of V_g_ and the among line variance of clonally related strains as an estimate of V_m_. We detected that V_g_/V_m_ is lowest in areas of largest plasticity around dorsal positions 100 to 250, in second instar animals exposed to predation risk (Fig. 3B). The localized reduction of V_g_/V_m_, coupled with the reduction of V_g_ in the induced state (Fig. 2F,G), provides strong evidence for stabilizing selection operating on plasticity.

## Discussion

We tested the hypothesis that a classic example of phenotypic plasticity - predator induced defense in *D. pulex* - is subject to stabilizing selection. We provide two lines of evidence to support our hypothesis: we show that standing genetic variation is reduced in regions of greatest plasticity (Fig. 2) and we show that standing genetic variation in plasticity is substantially lower than mutational variation (Fig. 3). Although the intuitive model that plasticity is directly related to fitness forms the basis for a century of work studying the anti-predator responses of *D. pulex* [28, 36] and other related Cladocera [37, 38], to our knowledge, this hypothesis has not been directly tested. Our work, therefore, provides novel insight into the evolutionary history of phenotypic plasticity.

Prior research has demonstrated that aspects of phenotypic plasticity can evolve in response to changes in the environment [3, 4]. Theoretical models that explain the evolution of plasticity generally assume that genetic diversity is sufficiently present [39, 40], and this assumption is generally realized when examining empirical data in a wide variety of species (e.g., [41]). Despite the ubiquity of genetic variation in plasticity across the tree of life, and genetically based phenotypic variation of fitness-related traits in general [22], determining the evolutionary forces that act on that variation remains a fundamental challenge [42]. Determining the evolutionary history of phenotypic variation requires a comparative approach, and often comparisons are made between populations to study local adaptation [8, 17] or across taxa to study diversification [43, 44]. As a consequence, identifying the forces that shape genetically based phenotypic variation within populations remains relatively understudied.

In order to gain insight into the evolutionary determinants of phenotypic variation within populations, it is critical to understand the extent of mutational variance. Mutational variance is generally considered deleterious [45], and therefore the rate at which it is removed from populations should reflect the strength of purifying selection [46]. Although directly observing the trajectory of new mutations is challenging, comparisons of standing genetic variation to mutational variation can yield insight into the expected persistence time of new, deleterious mutations [46], the mutational target size [47] and the genetic architecture of different classes of traits [34]. For instance, the values of V_g_/V_m_ that we observe are consistent with strong stabilizing selection which removes deleterious mutations quickly from the population via natural selection acting directly on the plasticity and not, in contrast, to a model of stabilizing selection generated via pleiotropy [32].

We detected values of V_g_/V_m_ in areas of largest plasticity are substantially lower than previously reported data (e.g., [45]). Thus, the question arises why mutational variance among clonally related strains is larger than the observed genetic variation among individuals from the outbred population in the area of greatest plasticity. Our analyses indicate that neither experimental artifacts nor maternal effects are the sole drivers of the observed substantial variation among clonally related strains (Fig. 3A) and the small values of V_g_/V_m_ (Fig. 3B). Consequently, we can only speculate on potential factors driving the observed pattern. The outbred population is derived from individuals that recently hatched from sexual ephippia deposited sometime in the past, whereas the clonally related strains were present in the population an extended period of time leading up to the point of collection. If the strength of predation varies through time, selection events further in time in the past may have depleted diversity in clones that result from sexual reproduction (i.e., our outbred population), while clonally strains may reflect more recent population history. Although the exact mechanism of the large among-line variance of the genetically similar clones is unclear, our analysis (Fig. 3) suggests that it is not solely generated by experimental artifact or maternal effects.

Examination of the evolutionary forces acting on plasticity is important for the interpretation of the evolutionary history of this population and also for assessing its evolutionary potential. While our data show that plasticity is subject to strong stabilizing selection, we also show that there is ample mutational variance for plasticity (Fig. 3). Mutational variance for plasticity could facilitate rapid adaptive evolution following shifts in the predator composition in the aquatic community due to climate change [48, 49] or other anthropogenic factors (e.g., [50]), and could thus be an important factor facilitating population persistence of *D. pulex* and other organisms, with consequences for ecosystem stability and function [51].

## Materials and Methods

### Study system

Our data come from a population of *Daphnia pulex* located in the Kilwood Coppice Nature Reserve in the Dorset region of the southern UK (grid reference: SY 93599 82555). Genotypes used in this study were sampled from two partly interconnected seasonal ponds with predominantly invertebrate predators during early spring in 2016 and 2017 (Suppl. Table 2). In the lab, sampled live individuals were established as iso-female clonal lineages and maintained in artificial hard water (ASTM; [52]) with seaweed extract (marinure; Wilfrid Smith Std., Northans, UK) under standard conditions: 15 animals L^-1^ ASTM were fed three times a week with *Chlorella vulgaris* (2×10^5^ cells ml^-1^; >1.5 mg carbon L^-1^) and reared under 16:8h light:dark conditions at 20°C.

### Genotyping

#### Sequencing

For DNA extractions, multiple adult *Daphnia* were placed into artificial hard water containing antibiotics (Streptomycin, Tetracycline, and Ampicillin, 50 mg L^-1^ of each) and fed Sephadex G-25 Superfine (cross-linked dextran gel) beads for 48 hours in order to minimize bacterial and algal contamination in downstream sequencing analyses. 5-10 individuals from each clonal lineage were then used for DNA extraction using Beckman-Coulter’s Agencourt DNAdvance kit. Individuals were homogenized using metal beads and a bead beater prior to DNA extraction. RNA was removed using RNase followed by an additional bead clean-up. DNA was quantified using the broad-range Quant-iT dsDNA kit (ThermoFisher Scientific) and an ABI plate reader and normalized to 1 or 2 ng ul^-1^ prior to library construction. Full genome libraries were constructed using a modified down Nextera protocol [53]. Libraries were size selected for fragments ranging from 450-550bp using Blue Pippin and quality checked using BioAnalyzer. Libraries were sequenced using the Illumina HiSeq 2500 platform.

#### Mapping, SNP calling, and SNP filtering

Nextera adaptor sequences were removed using Trimmomatic version 0.36 [54] and overlapping reads were merged using PEAR version 0.9.11 [55]. Assembled and unassembled reads were separately mapped to a European *D. pulex* reference genome ([31]) using bwa mem [56]. The entire reference genome was used for mapping, but only reads that mapped to *Daphnia* scaffolds, had quality scores greater than 20, and were primary alignments were used for further analysis. PCR duplicates were removed using Picard’s *MarkDuplicates* function [57]. GATK HaplotypeCaller (version 4.0, [58, 59]) was used to call SNPs. We removed SNPs that were within 10 base pairs of indels. SNPs were then hard filtered using GATK’s recommendations for organisms with no reference SNP panel (QD < 2, FS > 60, MQ < 40, MQRankSum < -12.5, and ReadPosRankSum < -8). Individual genotype calls with low quality scores (GQ < 10) were set as missing data.

#### Clonal assignment

Individual field isolates were assigned to clonal lineages based on patterns of identity by state (IBS). IBS was calculated using the snpgdsIBS function in SNPRelate [60], with a minor allele frequency cutoff of 0.001 and a missing rate of 0.15. We classified individual field isolates as coming from the same clonal lineage if pairwise identity by states was greater than 0.965 (see [31] for more details). We identified 50 genetically unique clonal groups, of which one cluster was represented by 56 genetically similar strains, yielding 105 strains that were used for phenotypic analysis. We investigated patterns of relatedness by calculating *IBS0* and kinship coefficients using the program KING [61]. KING was run using the “kinship” command with the input data filtered to include SNPs with a minor allele frequency cutoff of 0.05.

### Phenotyping

#### Experimental exposures

Phenotypic data were collected from 105 iso-female lines from the Killwood Nature Preserve (Suppl. Tab. 2). To establish predation risk conditions, we generated predator cues from frozen midge larvae following established protocols [62]. Homogenized midge larvae extracts were filtered, followed by solid-phase extraction using a C_18_ column (Agilent) to recover the active compounds that generate strong morphological responses in *D. pulex*. For experimental exposures, animals were kept under standard conditions for three generations. Subsequently, at least two gravid *Daphnia*, carrying embryos in E_3_ stage (∼18 hours before parturition; *sensu* [63]), were placed in individual jars containing 50 ml hard artificial pond water [52], algae (2×10^5^ cells ml^-1^ *Chlorella vulgaris*), liquid seaweed extract and 0 or 0.5 µl ml^-1^ *Chaoborus* predator cue concentrate. After parturition, two neonates were randomly selected from each of the two mothers per treatment (Suppl. Fig. 2) and placed individually in 50 ml glass vials containing the same medium as their maternal environment. For three to four consecutive days, each animal was photographed daily (Leica S8AP0 microscope; Leica EC4 camera) and subsequently transferred to a new glass vial containing fresh media and predator cues.

#### High-throughput image analysis

We assessed phenotypic changes using an automated image analysis pipeline hereafter referred to as *DAPCHA* (see Suppl. Methods). To validate accurate performance of *DAPCHA*, all images were manually checked, and if required, landmarks manually curated. We recorded manual estimates of units of microstage meter (>100 randomly chosen images) via two separate runs of estimation, and animal lengths and morphological induction (>700 randomly chosen images) via three independent observers using ImageJ software ([64]; Fiji plugin: [65]). We manually measured animal length from the tip of the head to the base of the tail. Morphological induction, based on the presence of a pedestal, was manually scored using a previously defined scoring system [28, 29]. Developed spikes in the nuchal area, later referred to as neckteeth, were counted individually.

### Instar assignment

Animals were retroactively assigned to distinct developmental stages (i.e., first and second instar) by fitting a mixture model on animal dorsal area and animal length, using the Mclust package for R [66].

### Analysis of variance

To assess the contribution of genotype, treatment, and their interaction, we fit a linear model (dorsal height ∼ genotype * treatment + batch), followed by Type II sums of squares implemented by the Anova() function in the car package for R [67] for significance testing, and estimation of effect sizes using the effectsize package for R [68].

### Magnitude and position of induced defense

To estimate how predation risk altered the magnitude and position of the morphological defense structures, we applied a phenotypic trajectory analysis [69] to the multivariate data matrix of dorsal height at each *i^th^* position along the dorsal edge using the geomorph package for R [70–72]. We fit a model where the response variable is the multivariate dorsal height x position matrix among genotypes versus treatment (control – predation). We estimated the overall impact of predation risk using 1000 permutations via the procD.lm() function. This was followed by assessment of the direction and magnitude statistics via the trajectory.analysis() function. Visualization of the morphology and details about the shift in height and position of maximal induction were made with the plotRefToTarget() function.

To assess modularity and identify whether the region where the morphological defense is induced is correlated (or not) with the rest of the body, we applied the modularity.test() function in the geomorph package for R [70–72]. We tested for the presence of modularity between areas of largest plasticity (i.e., head area) and the remaining dorsal areas along the carapace.

#### Correlation analysis

To assessed how much offspring resembled one another when released from (i) the same mother and the same clutch, (ii) the same strain but different mother, (iii) randomly drawn strains from the clonal assemblage of genetically related strains, and (iv) randomly drawn individuals from each experimental batch, we performed a correlation analysis for two key phenotypic traits, animal length and maximal dorsal height, using the R package robcor [73]. In order to estimate correlation coefficients for randomly drawn clone pairs from among genetically similar strains or batches, we performed 100 bootstraps. We also permuted phenotypic data in order to obtain a NULL distribution of correlation coefficients (n = 1000).

#### Heritability estimation

We estimated broad-sense heritability of phenotypic traits using the R package MCMCglmm [74]: dorsal height ∼ 1, ∼ clone + batch. Models were fitted with a burn-in of 15,000 and sampling that produced 1300 estimates of the posterior distribution from 65,000 iterations of the chain. All models were checked for autocorrelation in the chains. We calculated V_g_/V_m_ as mean(log_10_(posterior distribution _standing variation_) - log_10_(posterior distribution _mutational variation_)).

We next estimated narrow-sense heritability of phenotypic traits in the genetically divergent clones using GCTA [75]. We generated a set of genome-wide representative SNPs, and then calculated a genetic relatedness matrix from these SNPs with the --make-grm flag in GCTA. For heritability estimations, we used the flags --reml, --reml-alg 1, accounting for *batch* as covariate. In order to estimate heritability estimates for random data (i.e., NULL distribution), we permuted genome identifiers before calculating genetic relatedness matrices.

### Statistical analysis and plotting

All analyses were performed using R version 3.5 [76]. The following packages were used for general analysis and plotting: *ggplot2* [77], *cowplot* [78], *data.table* [79], f*oreach* [80], *doMC* [81], *ggbeeswarm* [82], and *viridis* [83].

### Data Availability

All scripts and code used for data analysis and plotting are available at https://github.com/beckerdoerthe/SelectionPlasticity. DAPCHA is available at https://github.com/beckerdoerthe/Dapcha_v.1. All raw images and processed data used to generate figures are deposited in Zenodo (DOI 10.5281/zenodo.4738526). All sequencing reads are available from the Sequence Read Archive (PRJNA725506).

## Acknowledgments

AOB was supported by the National Institutes of Health (R35 GM119686) and by start-up funds provided by the University of Virginia. This project has received funding from the European Union’s Horizon 2020 research and innovation programme under the Marie Sklodowska-Curie grant agreement No 841419. The authors acknowledge Research Computing at The University of Virginia for providing computational resources and technical support that have contributed to the results reported within this publication (https://rc.virginia.edu). The authors would also like to thank the Dorset Wildlife Trust for granting access to the field site.

## Author Contributions

<u>DB</u>: Conceptualization, Data curation, Formal analysis, Investigation, Methodology, Resources, Software, Visualization, Writing - original draft, Writing - review & editing; <u>KBK</u>: Investigation, Resources, Writing - review & editing; <u>RP</u>: Investigation, Writing - review & editing; <u>AE</u>: Investigation, Writing - review & editing; <u>EV</u>: Investigation, Writing - review & editing; <u>APB</u>: Conceptualization, Resources, Writing - review & editing; <u>AOB</u>: Conceptualization, Data curation, Formal analysis, Funding acquisition, Investigation, Methodology, Project administration, Resources, Software, Supervision, Visualization, Writing - original draft, Writing - review & editing.

## Competing Interest Statement

All authors declare no competing interests.

## Supplemental Information

**Supplemental Table 1.**
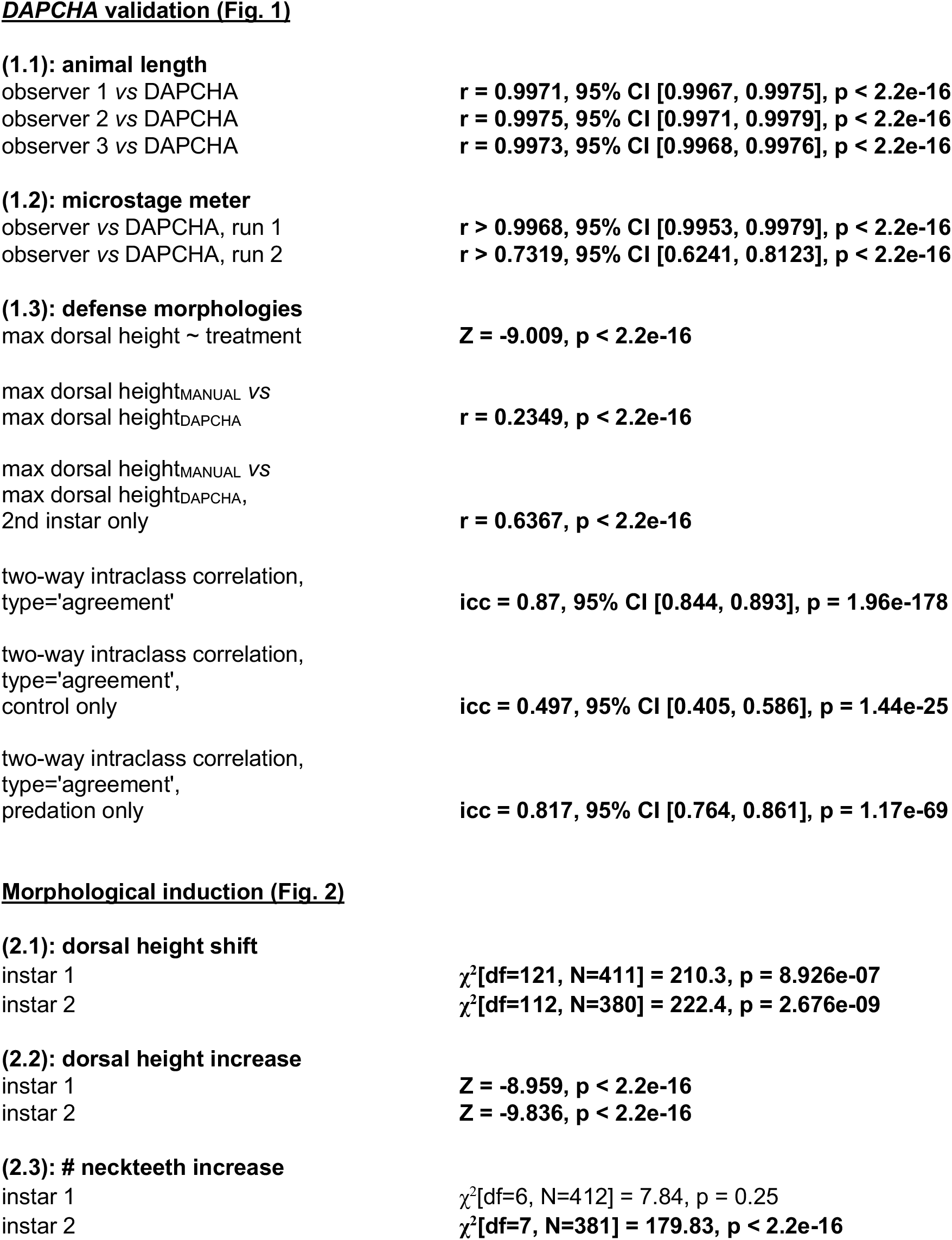
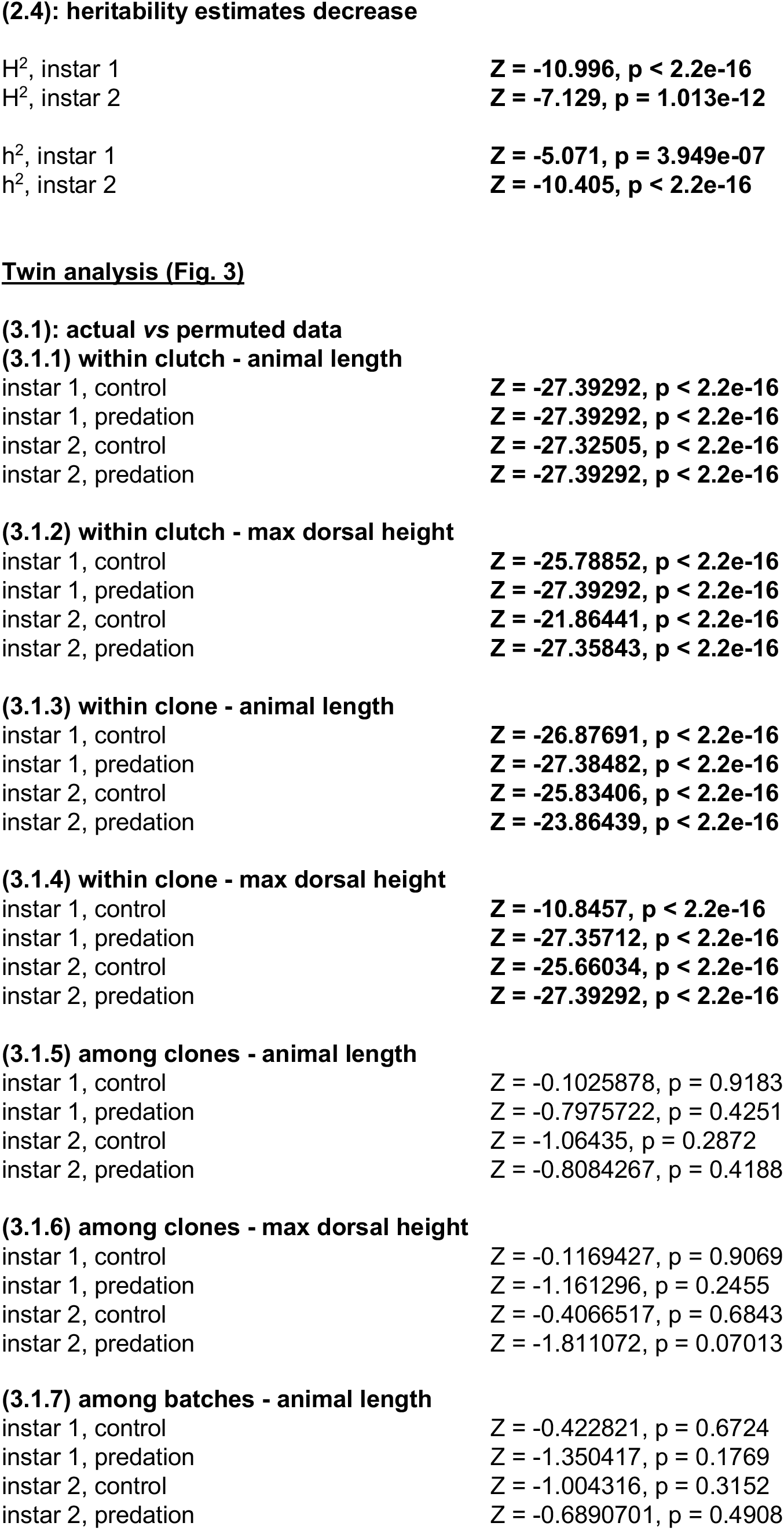
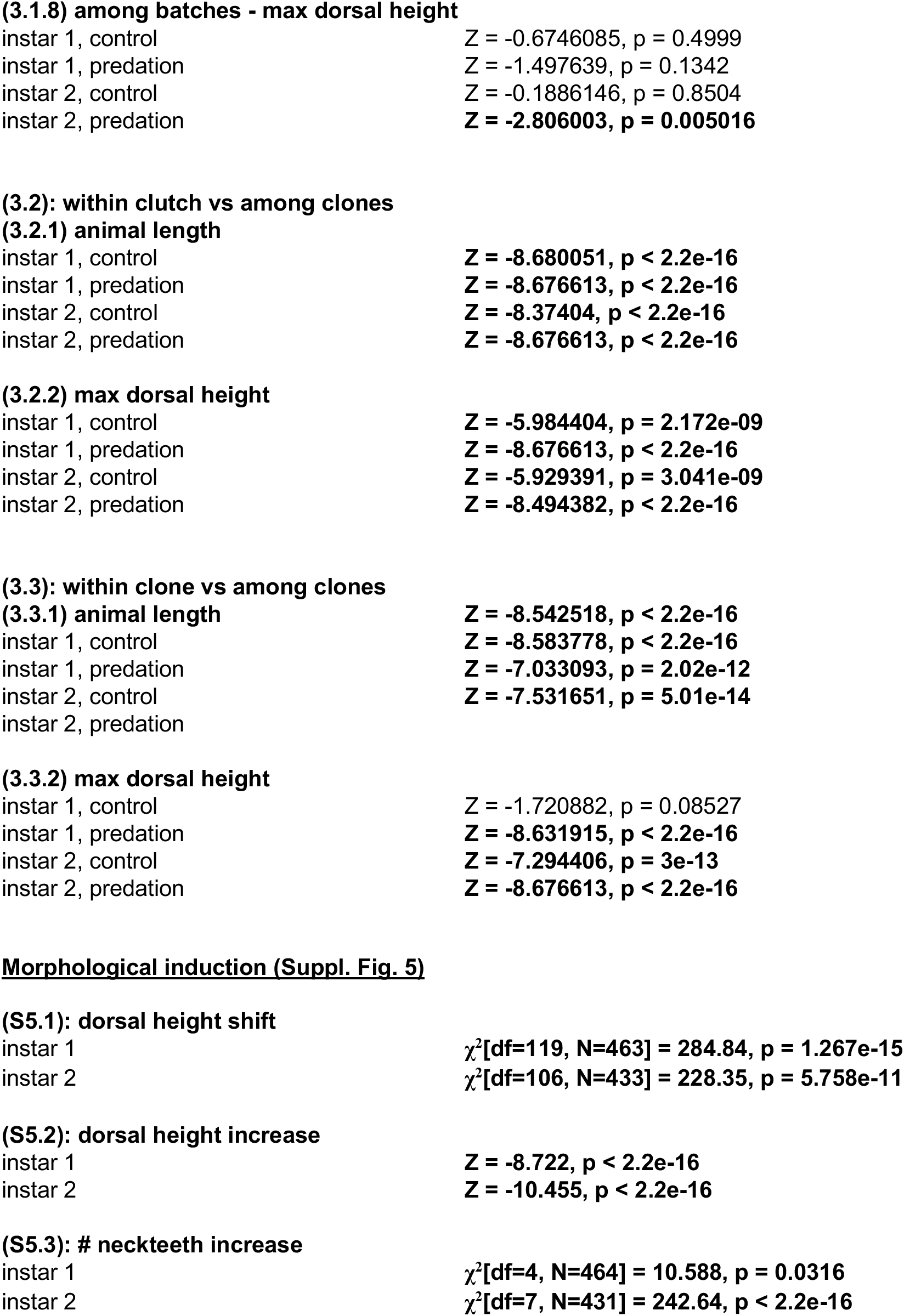
List of statistical tests outcomes.

**Supplemental Table 2.**
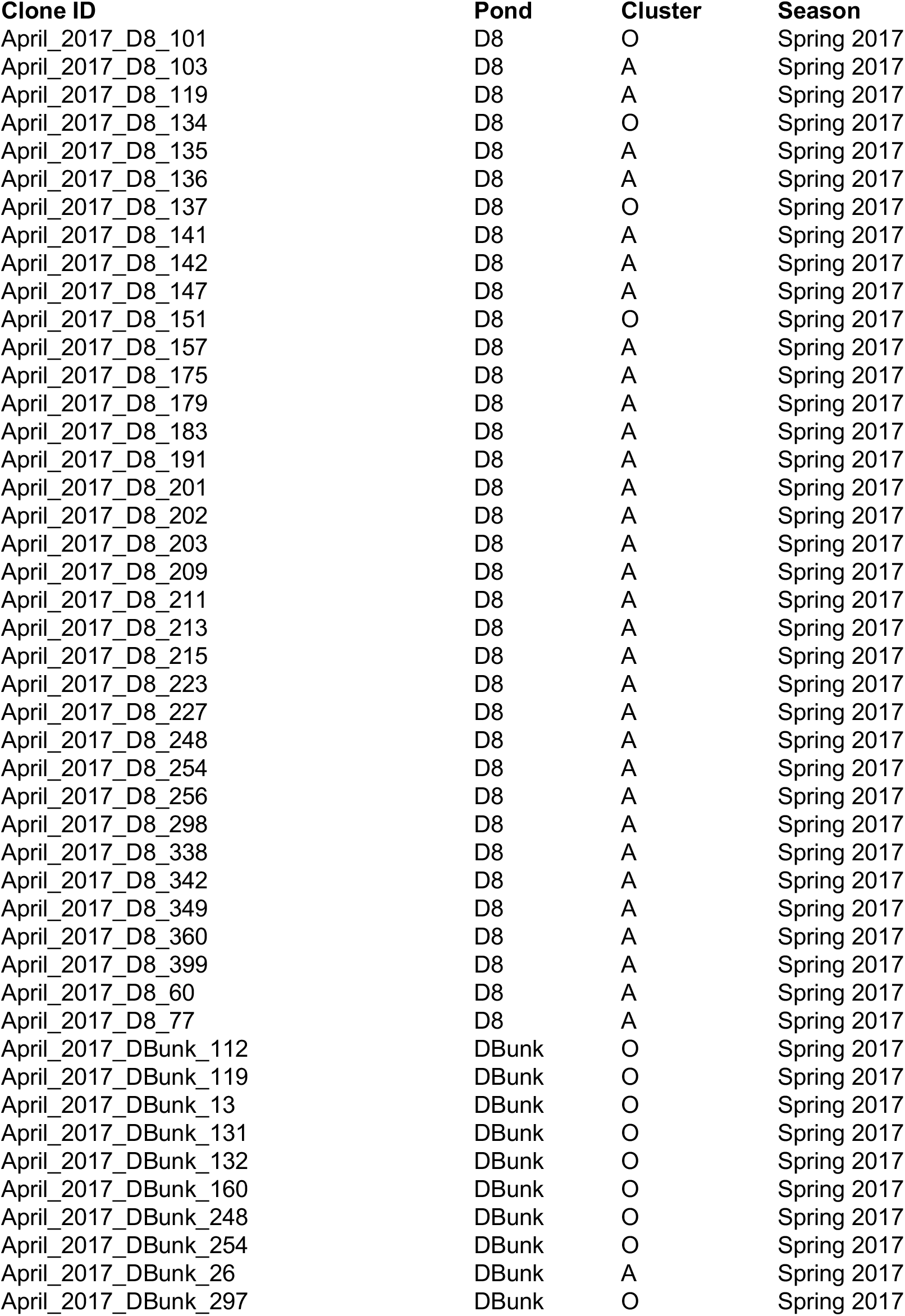
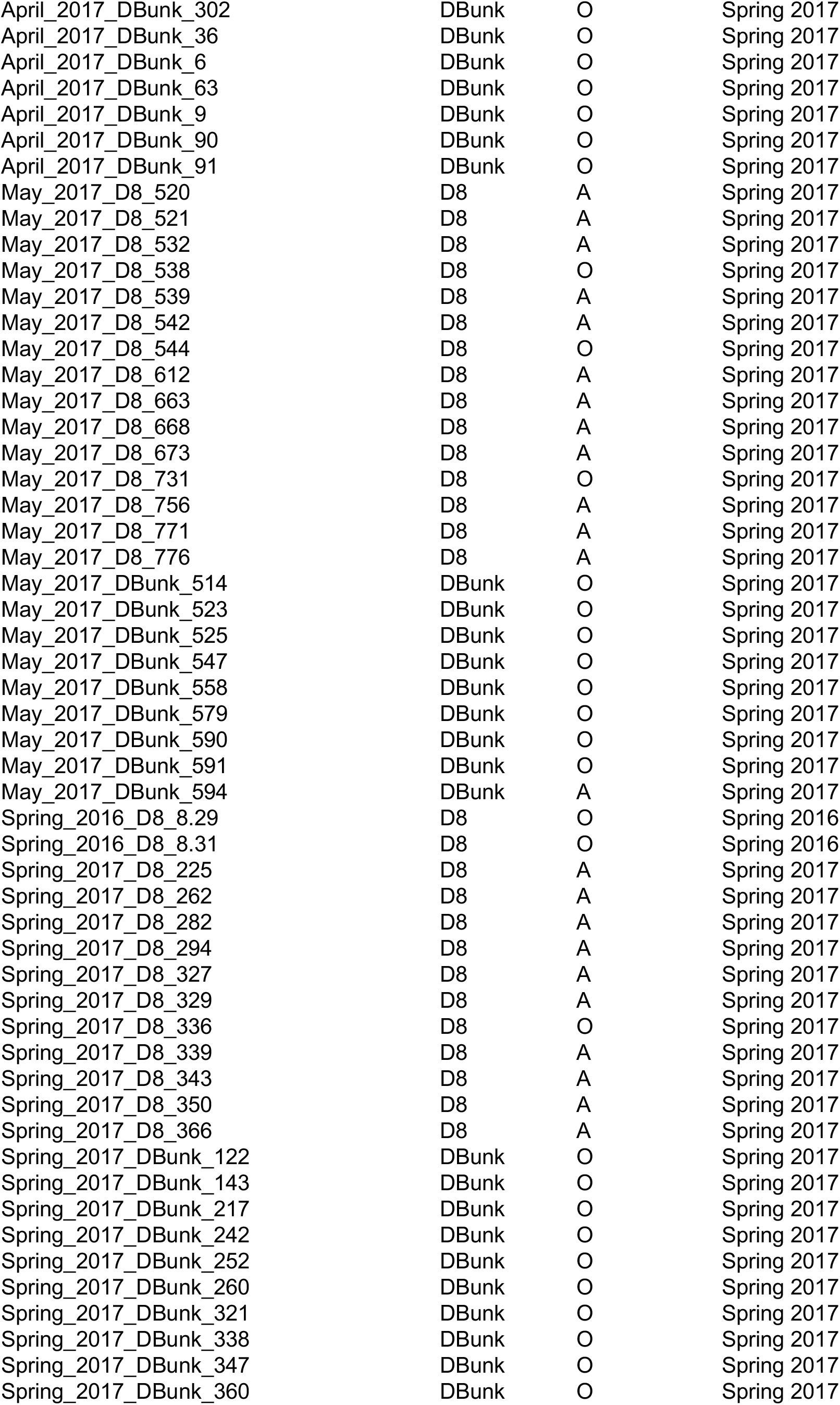
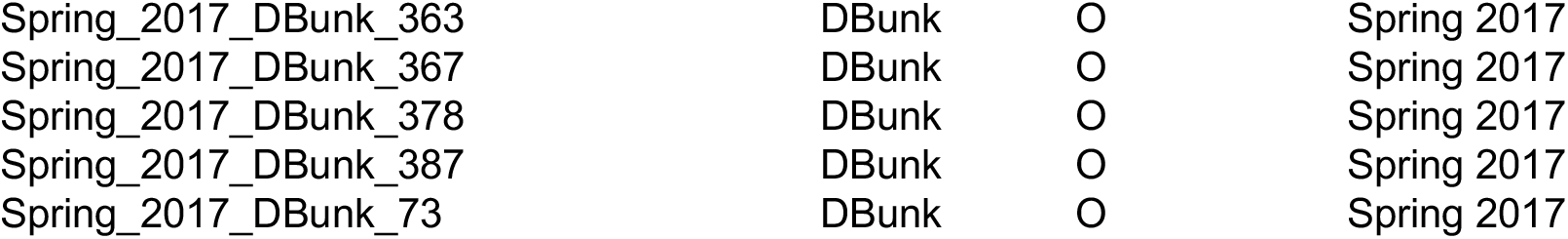
List of *Daphnia pulex* clones used in this study, including information on pond identity and sampling season. Cluster information indicates whether strains are genetically similar (‘cluster A’) or genetically unique (‘cluster O’).

## Supplemental Figures

**Supplemental Figure 1.**
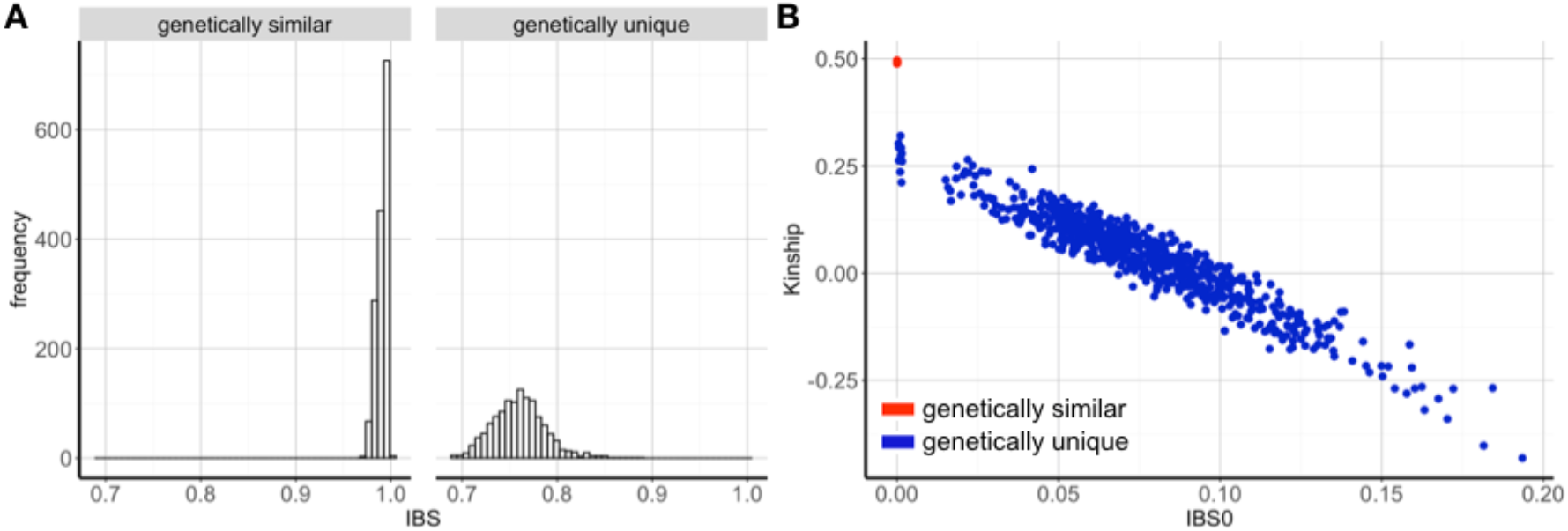
Genetic diversity among genetically similar and unique strains. (A) Distribution of pairwise IBS values between all genetically similar (left) and genetically unique (right) strains. (B) Relationship between IBS0 and kinship as calculated in the program King for pairwise combinations of individuals genotyped from the sampled population. Red and blue circles depict genetically similar and unique strains, respectively. Note that in (B) all comparisons between genetically similar strains are stacked on top of each other.

**Supplemental Figure 2.**
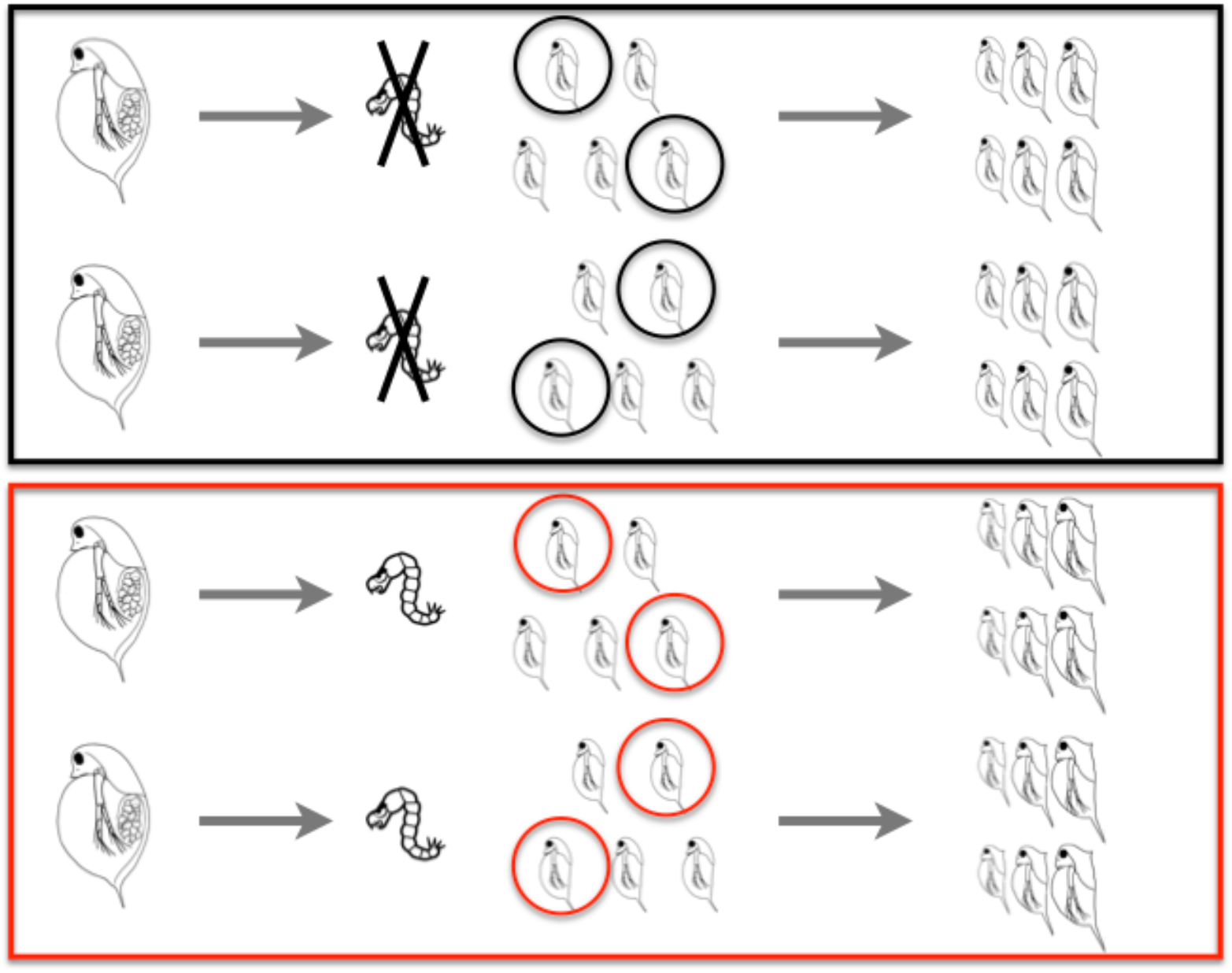
Experimental design. For experimental exposures, two mature *Daphnia pulex* carrying embryos in E3 stage (∼18 hours before parturition; *sensu* [63]) were placed in individual jars containing medium with (red box) and without (black box) predator cues. After parturition, two neonates were randomly selected from each of the two mothers and placed in individual vials containing the same medium as their maternal environment. Subsequently, animals were monitored for 3-4 consecutive days, with daily photographs taken.

**Supplemental Figure 3.**
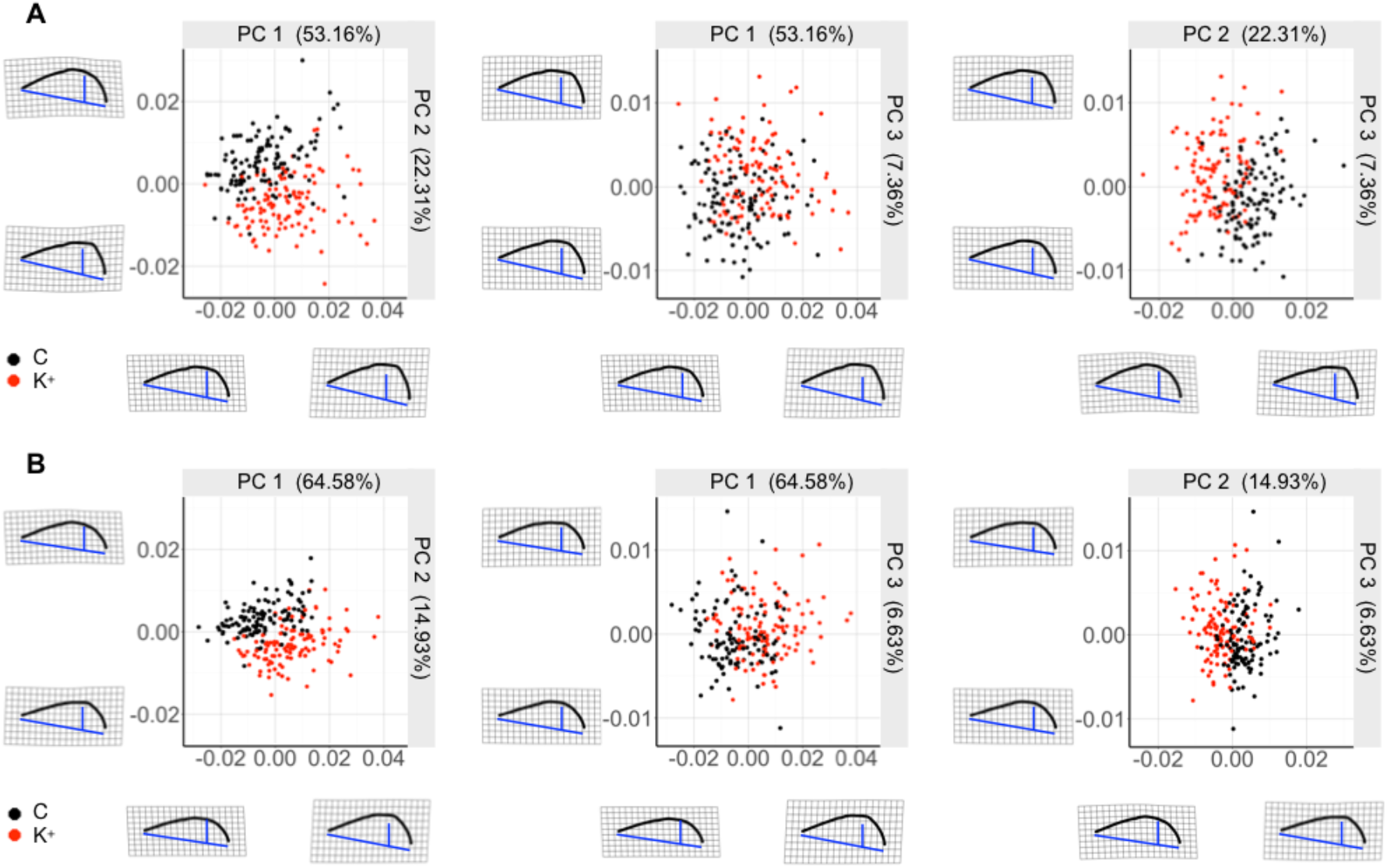
*Chaoborus* induced shape variation in *D. pulex*. Visualization of the first three main axes of dorsal shape variation in first (A) and second instar (B) *Daphnia* using principal component (PC) analysis of procrustes data. Colors indicate treatment conditions (control: black points, predation: red points). Warp-shape diagrams highlight distinctive patterns of shape variations along the principal components: PC1 represents shape differences in dorsal height, while PC2 and PC3 characterize the development of predator induced defense morphologies and shifts in their dorsal position, respectively.

**Supplemental Figure 4.**
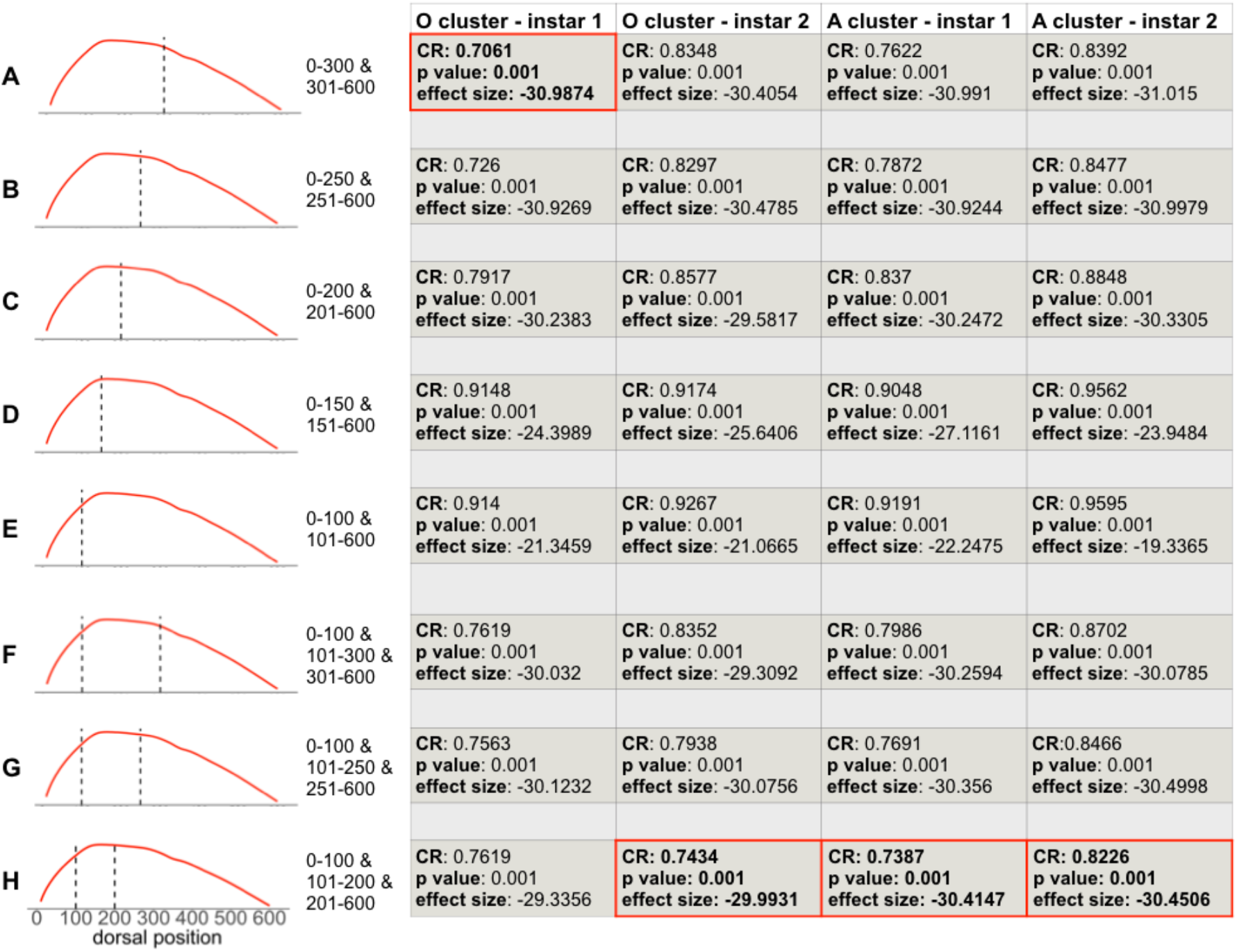
Modularity of predator induced defenses along dorsal axis. A formal modularity analysis, testing for the presence of distinct phenotypic modules along the dorsal axis, indicates that plastic responses in the nuchal area are independent of changes in other body parts: there is strong statistical evidence for either two (A-E) or three (F-H) independent dorsal modules separating areas of largest plasticity (i.e., head area) and the remaining dorsal areas along the carapace. The extent of modularity is described by a CR coefficient in proposed modules compared against randomly assigned groups of landmarks using a random permutation procedure. O cluster: genetically unique strains; A cluster: genetically similar strains.

**Supplemental Figure 5.**
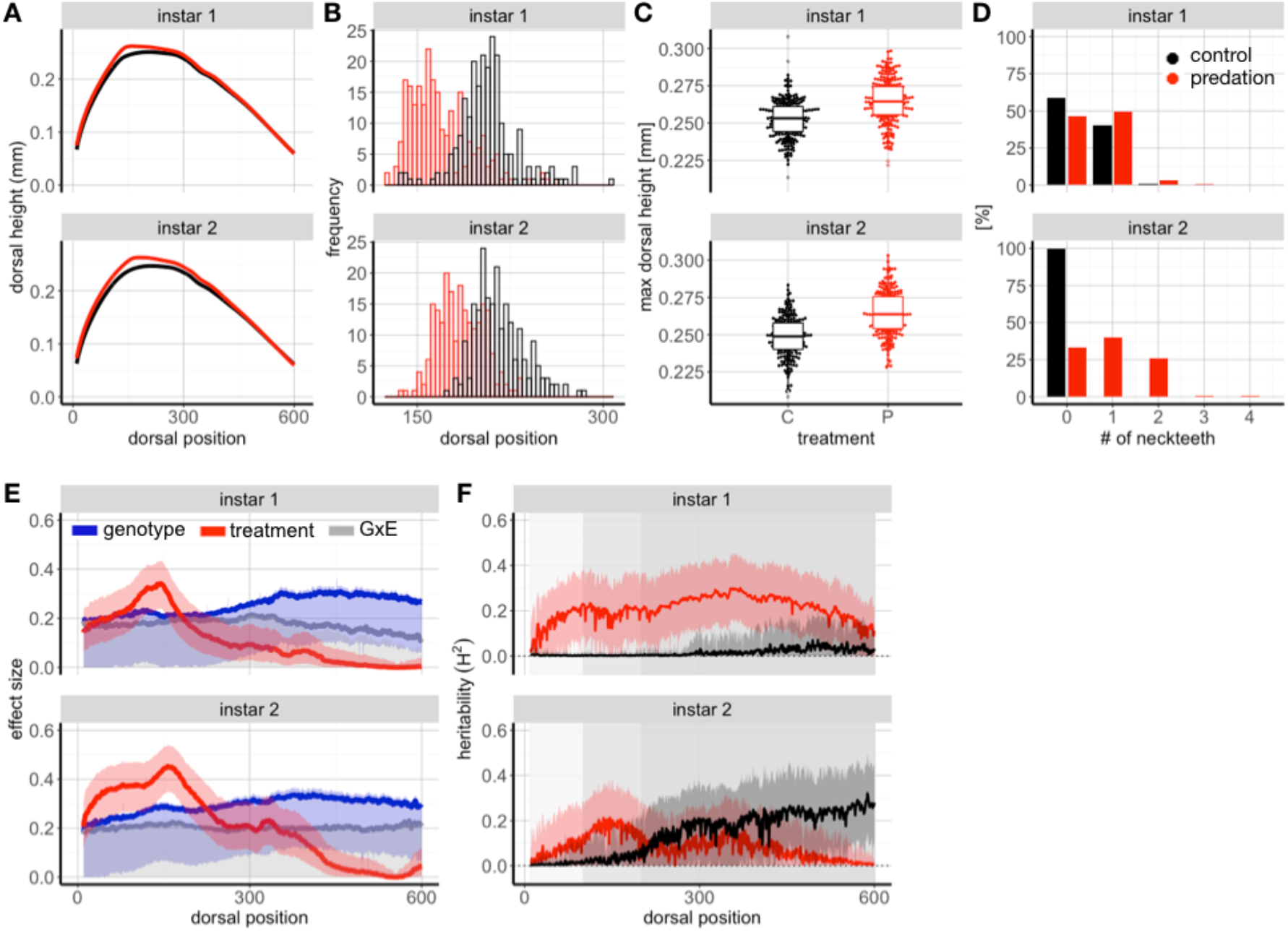
Effects of predation risk on morphological changes in genetically similar strains. (A) Risk of predation induces plastic responses, with strongest phenotypic changes observed in the nuchal area (around dorsal positions 100-250). (B,C) In response to predation, maximum dorsal height increases and shift towards anterior head regions. (D) In addition, the number of neckteeth increases und predation risk. Notably, variation in morphological changes within genetically identical clones is as pronounced as that observed among genetically unique clones (see Fig. 2). (E) Effect sizes from analyses of variance along the dorsal shape reveal distinctive patterns of treatment (i.e., predation risk, red line), genotype (blue line), and GxE (grey line) effects on morphological changes. Shaded areas indicate upper (0.95) and lower (0.05) confidence intervals. (F) Broad sense heritability estimates of dorsal height vary along the dorsal axis in response to control conditions (black line) and predation risk (red line) in genetically similar clones. Grey rectangles highlight morphological independent shape modules, separating head and posterior body areas (see Suppl. Fig. 4).

## Supplemental Methods

### High-throughput image analysis

We assessed phenotypic changes using an automated image analysis pipeline hereafter referred to as *DAPCHA*, using Python version 2.7. First, we identified the eye as a common landmark for each individual image: we first identified the center of the eye by calculating the mean of the darkest 2.5% of all pixels in the image. The rest of the pixels comprising the eye were identified using a 4-way flood-fill style algorithm on the image after filtering with a Canny edge detector implemented in OpenCV [84], Gaussian blur sigma = 0.5, Canny edge filter with lower threshold = 0, and higher threshold = 50). Next, we identified the center pixel of the animal by taking the mean of all edge pixels produced by a Canny edge filter (lower threshold = 0, higher threshold = 50) applied to a Gaussian blurred image (sigma = 1.25). Then, the tip of the tail was approximated by computing the dot product between the *animal center - eye center vector* and every *animal center - edge pixel vector*. The maximum dot product of these were considered the tail tip, since the tail tip should lie the furthest away from the eye. To estimate the animal position within the image, an ellipse was fitted to the edge pixels, providing the center, major and minor axes and orientation of the body of the animal. The major axis vertices were assigned anatomical directions (anterior and posterior) by comparing their locations relative to the eye center. Similarly, the minor axis vertices locations were compared to the tail tip. The ventral and dorsal-most eye points were found by translating the minor axes vectors to originate at the center of the eye and finding the shortest distance between all eye points and the terminal point of the vector. Next, the base of the tail was estimated by finding where the tail spine begins to thicken and joins the bottom of the main carapace. A line segment was drawn between the tail tip towards the eye and divided into 100 points. Starting at the tail tip, a second line segment was drawn between each of the 100 points and a point perpendicular to the eye-tail line. Along this line, the outermost edge pixel was identified (i.e., the edge of the tail spine). If this pixel was above a distance threshold (1/5^th^ of the distance between the starting and ending points of the line segment), the search was terminated, and the pixel was designated as the *tail point*. The line segment with this tail point edge pixel was extended dorsally, and the furthest edge pixel on that side was designated as the *dorsal tail point* (i.e., the base of the tail on the dorsal side of the animal). To locate the head, a line segment was drawn between the new tail point and the dorsal eye point. This line segment was then extended until no more edge pixels were found. The last edge pixel was designated as the *head point*. Using these landmarks, we calculated animal length, animal area, eye area, tail spine length.

After landmark detection, the dorsal edge of the carapace was traced by connecting edge pixels between checkpoints along the dorsal edge. These checkpoints were determined by drawing a line segment between the eye and tail, dividing this segment into ten points, and then finding the most dorsal edge pixel along the line perpendicular to the eye-tail line containing these ten points. Due to noise and variability, some of these checkpoints may fall far from the actual dorsal edge of the carapace. These wayward points were pruned from the checkpoint list by calculating the distance of checkpoint from the line drawn between its neighboring checkpoints, discarding it if too far away from the line. After checkpoints were determined, the dorsal edge was traced in segments between checkpoints. Via an iterative search process, one checkpoint per each segment was designated as the *target*, and one as the *current point*. Relative to the *current pixel*, the next pixel should be the one that was both dorsally and posteriorly located and further away from previously traced points. To calculate a position score, all of the edge pixels around the current point within a small neighborhood were indexed and a score was calculated by adding the dot product between the index and three different vectors: a vector between the *current point* and the *target point*, a vector between the last *current point* and the *current point*, and the vector starting at the animal center and ending at the *current point*. The edge pixel corresponding the maximum score was labeled as a dorsal edge and then set as the *current point*. This procedure continued until the *target point* was reached. Finally, all of the dorsal edge points from each segment were concatenated together. In order to account for differences in number of pixels due to size variability among animals, we equalized the number of points between dorsal edge traces: we implemented a greedy algorithm using linear interpolation to add points to the dorsal edge until a set target number of points was reached at 700. Dorsal edge detection allowed estimation of morphological changes along the entire dorsal shape axis, later referred to as *dorsal height*.

### Pixel-to-mm extraction from micrometer images

Each slide micrometer image was cropped to include only the scale line. This was done by automatically detecting the dark edges of the micrometer ring and cropping them out. Next, we applied contrast limited adaptive histogram equalization [84] to increase the contrast of the scale against the brightfield background. We then applied a Canny edge filter [84] (lower threshold=0, higher threshold=175) to the high contrast image and used a Hough [84] line detector to find the largest line in the image (i.e., the cross bar in the scale line). Next, we extracted the intensity of the image along a line parallel to the cross bar, drawn along the tick marks in the scale line. Finally, we applied a Fourier transform to the coordinates of the intensity peaks along this line to measure the frequency of the tick marks. This frequency corresponds to the inverse pixel-to-mm ratio divided by a factor of 40.

## Notes

### Competing Interest Statement

The authors have declared no competing interest.

